# Real-time Ratiometric Imaging of Micelles Assembly State in a Microfluidic Cancer-on-a-chip

**DOI:** 10.1101/2020.03.08.978783

**Authors:** Natalia Feiner-Gracia, Adrianna Glinkowska Mares, Marina Buzhor, Romen Rodriguez-Trujillo, Josep Samitier, Roey J. Amir, Silvia Pujals, Lorenzo Albertazzi

**Affiliations:** Institute for Bioengineering of Catalonia, The Barcelona Institute of Science and Technology (BIST), Carrer Baldiri Reixac 15-21, 08024 Barcelona, Spain; Department of Biomedical Engineering, Institute for Complex Molecular Systems (ICMS), Eindhoven University of Technology, 5612AZ Eindhoven, The Netherlands; Department of Organic Chemistry, School of Chemistry, Faculty of Exact Sciences, Tel-Aviv University, Tel-Aviv 6997801, Israel; Tel Aviv University Center for Nanoscience and Nanotechnology, Tel-Aviv University, Tel-Aviv 6997801, Israel; BLAVATNIK CENTER for Drug Discovery, Tel-Aviv University, Tel-Aviv 6997801, Israel; Department of Electronic and Biomedical Engineering, Faculty of Physics, University of Barcelona, Carrer Martí I Franquès 1, 08028 Barcelona, Spain; Networking Biomedical Research Center in Bioengineering, Biomaterials and Nanomedicine (CIBER-BBN), 28029 Madrid, Spain

**Keywords:** Micelle, Stability, Cancer-on-a-chip, supramolecular, nanoparticle, microfluidic

## Abstract

The performance of supramolecular nanocarriers as drug delivery systems depends on their stability in the complex and dynamic biological media. After administration, nanocarriers are challenged by confronting different barriers such as shear stress and proteins present in blood, endothelial wall, extracellular matrix and eventually cancer cell membranes. While early disassembly will result in a premature drug release, extreme stability of the nanocarriers can lead to poor drug release and low efficiency. Therefore, comprehensive understanding of the stability and assembly state of supramolecular carriers in each stage of delivery is a key factor for the rational design of these systems. One of the key challenges is that current 2D *in vitro* models do not provide exhaustive information, as they do not fully recapitulate the 3D tumor microenvironment. This deficiency of the 2D models complexity is the main reason for the differences observed *in vivo* when testing the performance of supramolecular nanocarriers. Herein, we present a real-time monitoring study of self-assembled micelles stability and extravasation, combining spectral confocal microscopy and a microfluidic tumor-on-a-chip. The combination of advanced imaging and a reliable organ-on-a-chip model allow us to track micelle disassembly by following the spectral properties of the amphiphiles in space and time during the crucial steps of drug delivery. The spectrally active micelles were introduced under flow and their position and conformation followed during the crossing of barriers by spectral imaging, revealing the interplay between carrier structure, micellar stability and extravasation. Integrating the ability of the micelles to change their fluorescent properties when disassembled, spectral confocal imaging and 3D microfluidic tumor blood vessel-on-a-chip, resulted in the establishment of a robust testing platform, suitable for real-time imaging and evaluation of supramolecular drug delivery carrier’s stability.

## Introduction

Supramolecular nanocarriers, such as liposomes or micelles, are broadly investigated as potential drug delivery systems (DDS) for cancer therapy.[1] [2] Since Doxil, a liposome-encapsulated Doxorubicin formulation, was approved in 1995,[3] the therapeutic efficacy of many nanosystems with different chemical features have been tested. However, low accumulation of the nanoparticles (NPs) in the solid tumor is still a major issue[4] and one of the key reasons for failures in clinical trials. One of the main challenges with the use of supramolecular nanocarriers is to control their dynamic nature, *i.e.* their assembly-disassembly equilibrium, which can determine *in vivo* success. NPs have to withstand the exposure to diverse environments affecting their properties, as a result the number of NP that arrive fully assembled to the target site can decrease by 500-fold.[5] Therefore, it is important to design nanosystems that are stable once injected into the body, avoiding premature disassembly and drug release and are “smart” enough to free the cargo when the target is reached.

Several factors can be responsible for the disassembly of supramolecular nanocarriers such as micelles. Right at the injection site, the NPs face a high dilution in blood and interactions with serum proteins, which may favor destabilization.[6] Moreover, the directed continuous flow present in the vessels induces shear stress, which may also cause the disassembly.[7] These events may result in premature drug release into the blood stream, causing undesired systemic cytotoxicity or uncontrolled drug distribution.[8] The next challenge comes from the selective extravasation through the layer of endothelial cells (ECs), lining the blood vessel’s internal wall. ECs, connected by characteristic tight junctions, create a physical barrier, which allows the diffusion of only small molecules.[9] Interactions with ECs membrane during extravasation, can further compromise micelle stability, which could cause the premature release of the cargo in the endothelial cells instead of inside the tumor cells. Nowadays, most of nanoparticles-based drug delivery systems rely on the enhanced permeability and retention effect (EPR), which is a phenomenon occurring in the tumor vasculature.[10] During EPR the tight endothelial junctions are impaired, creating gaps that allow larger molecules to leave the systemic circulation,[11] thus opening an escape route for the NPs.[12] Particles able to extravasate through the “leaky” endothelial barrier (EB), arrive to the extracellular matrix (ECM) surrounding the cancer cells. The ECM can differ in pH and shear stress in comparison with the blood vessel, moreover, NPs interactions with ECM components (collagen, glycosaminoglycan, proteoglycans) challenge their stability once more.[5] Eventually, nanocarriers arriving to the target site should be able to penetrate the tumor cells and release the drug once internalized. The predictability and understanding of the stability of supramolecular assemblies within these changing conditions, can pave the way to an improved design of nanosystems to be translated into the clinics. Therefore, a comprehensive study of NPs behavior in complex and dynamic environment requires a reliable, precise and versatile method. Here we address this challenge by combining spectral imaging and an organ-on-a-chip platform to study the stability of spectrally active polymeric micelles with highly tunable degree of amphiphilicity.

Recently, a Förster Resonance Energy Transfer (FRET) based approach has been used to follow the disassembly of supramolecular assemblies in blood serum,[13]–[16] and in animal models,[17] by encapsulation or covalent attachment of a FRET fluorophore pair into one micelle. In a recent work we employed PEG-Dendron amphiphiles that were functionalized with a spectral-shifting coumarin to track the stability of micelles in serum and cells. These micelles can self-report their disassembly by changing their fluorescent spectrum back to the intrinsic emission of the labeling coumarin dyes. In all cases, the change in assembly can be detected in complex media using spectral imaging, enabling to study the stability of the tested NP in space and time.

In this work we propose to combine the use of spectral imaging with microfluidic organ-on-a-chip technology to evolve conventional 2D studies into more realistic 3D models.[18],[19] By investigating 3D arrangement and transport phenomena in blood vessel and tumor microenvironment we aim to provide additional screening before use of animal models.[20] [21] [22] In the last years, researches have proposed versatile platforms to study cancer cells migration, [23] [24] [25] [26] vascular angiogenesis [27] [28] EB permeability^23 24^ and tumor penetration.^25 26^ Recently, more complex healthy and tumor blood vessel-on-a-chip have been designed in order to assess the differences in nanoparticles permeability in various conditions,^23 24 27^ demonstrating the appearance of increased endothelial permeability when cancer cells were present.[30] However, all microfluidics models developed up to now, were used for studying only the end time point of nanoparticles incubation, lacking time-resolved monitoring of their performance. We believe that it is very important to close the gap in understanding the crucial changes occurring in the supramolecular nanocarriers throughout their journey.

Herein, we combined continuous real-time monitoring of spectrally active polymeric micelles under flow using ratiometric confocal microscopy imaging, with 3D cancer blood vessel-on-a-chip model. The major aim of our work was to study the stability of different micelles in the various barriers and screen for the best performing candidate. The designed cancer-on-a-chip setup with our mapping method opens the way towards a new testing tool for optimizing the pre-clinical screening of nanoparticles.

## Materials and methods

### Microfluidic device

Microfluidic 3D culture chip DAX-1 (AIM Biotech) was used as a platform to reconstruct tumor microenvironment. LUC-1 connectors (AIM Biotech) were used to connect the chip inlets with luer connector ended PTFE tubing. The other end of the tubbing was connected to a syringe placed in a double syringe pump (Nexus Fusion 200) and filled with HUVEC (EndoGRO, Millipore) basal medium, used to constantly perfuse the chip for 48 – 72h.

### Cells and reagents

Human Umbilical Vein Endothelial Cells (Promocell) were used to recreate blood vessel lining and HeLa cells were used in to create tumor spheroids. HUVECs were cultured in EndoGRO Basal Medium (Millipore) supplemented with SCME001 kit (EndoGRO-LS Supplement 0.2%, rh EGF 5 ng/mL, Ascorbic Acid 50 μg/mL, L-Glutamine 10 mM, Hydrocortisone Hemisuccinate 1 μg/mL, Heparin Sulfate 0.75 U/mL, FBS 2%) and penicillin/streptomycin 1% (Biowest). HeLa cells were cultured in Dulbecco’s Modified Eagle Medium (DMEM, as received with L-Glutamine, 4.5 g/L D-glucose and pyruvate, Gibco) supplemented with FBS 5% (Gibco) and penicillin/streptomycin 1% (Biowest). HUVEC were cultured in 75 cm2 flasks and HeLa in 25 cm2 flasks at 37°C and 5% CO2. Cells were harvested using trypsin-EDTA (0.25%, Gibco) when reached 70-80% confluence.

### Cell culture in the Microfluidic device

Collagen gel at concentration of 2.5-3.0 mg/mL was prepared, introduced and polymerized according to the general protocol v5.3 (AIM Biotech). In brief, Rat tail collagen Type I (Corning Life Science) was mixed on ice with 10x PBS (Sigma Aldrich) and Phenol Red (Sigma Aldrich), and pH of the mixture was adjusted between 7-8 using 0.5 M NaOH (NaOH in pellets PanReac dissolved in MiliQ water), final volume was adjusted with MiliQ water (for healthy model) or suspension of HeLa clusters (for cancer model).

For preparation of cancer model microfluidic chip HeLa cells were seeded into 96-well ultralow attachment plate (Corning) at 0.5 – 1.5 k cells/well and cultured for 48-96h. Formed cell spheroids were harvested, centrifuged and resuspended in previously prepared collagen gel, resulting in few clusters (30-200 μm) per 10 μL of the gel. Prepared collagen was inserted into the central channel of 3D culture chip and allowed to polymerize during 30 min at 37°C and 5% CO_2_. After gel polymerization one of the lateral channels was prepared for HUVECs culture, by coating the channel with 50 μg/mL fibronectin (FN) from bovine plasma (Sigma Aldrich) during about 2h at 37°C. Remaining lateral channel was filled in with DMEM (HeLa culture medium) and closed using luer caps.

After the incubation time, the FN was washed away using 1x PBS (Gibco) and EndoGRO HUVEC medium. HUVECs were seeded to the prepared lateral channel at a density of 2.5-3.5M cells/mL. The 3D culture chip was flipped upside down to allow cell adhesion to the upper plane during 1.5-2.5h at 37°C and 5% CO_2_. Second batch of HUVECs cultured in another flasks was harvested and introduced to the same lateral channel at the same concentration as previously. The cells were then incubated for minimum 2h at 37°C and 5% CO_2_ in the upright placed chip to allow their attachment to the lower plane. Next, the chip was perfused with EndoGRO HUVEC medium at a flow rate 3-5 μL/min during 48 - 72 hours (as described above), until HUVECs reached confluency.

### Hybrids perfusion setup

Hybrids were prepared at a concentration of 480 μM in filtered PBS, sonicated for 5 minutes and let to equilibrate for at least 10 minutes. Prior to hybrid flowing into the chip they were mixed with EndroGRO HUVEC medium resulting in final concentration of 160 μM.

The microfluidic chip was placed into the on-stage incubator of a Zeiss LSM 800 Confocal microscope at a temperature of 37°C and 5% CO_2_, and connected to peristaltic pump (Ismatec, Reglo Digital, ISM597) with a silicon tubing (Tygon, Kinesis) to perfuse hybrids at real-time, at 15 uL/min, while imaging. Hybrids were excited using 405 nm laser and emission spectra was collected using two different PMT detectors to detect both monomer and micelle separately and simultaneous. The windows of detection were set as following: i) monomer 446-500 nm and ii) micelle 500-700 nm. Ratiometric images were obtained from dividing the micelle image by the monomer image, after applying a mask to each image were noise was removed.

To calculate the amount of hybrid able to extravasate we first sum up the signal of both windows. Next, we calculated the mean intensity signal of the vessel channel and used this value as the maximum concentration. Next the mean intensity signal of the gel channel was calculated and divided by the maximum signal concentration.

### Dextran perfusion

10kDa Dextran labelled with AlexaFluor568 (Thermo Fisher Scientific) was diluted in HUVEC (EndoGRO) medium at a final concentration of 1 μg/mL. The solution was perfused into the “blood vessel” of the chip at a flow rate of 5 μL/min using a syringe pump. The perfusion of dextran was monitored using Nikon Eclipse Ti2 epifluorescent microscope. The chip was placed in the on-stage incubator (Okolab) at a temperature of 37°C and 5% CO_2_, the perfused fluorophore was excited at 525 nm and emission collected at 650 nm.

### HeLa spheroid viability assay

The viability of HeLa cells within the spheroids were evaluated using Calcein (Fluka, Sigma Aldrich) and Propidium Iodide (Sigma Aldrich) to stain live and dead cells, respectively. First, cells were incubated with 10 μM Calcein solution for 20 min. at 37°C and 5% CO_2_ and then. Next, the cells were incubated with 10 μg/mL Propidium Iodide solution for 5 min. at 37°C and 5% CO_2_ and then washed with 1x PBS (Sigma Aldrich). The imaging was performed using Zeiss LSM 800 Confocal microscope. The Calcein and Propidium Iodide stained spheroids were excited at laser wavelength of 488 nm and 561 nm respectively and detection windows set at 400 – 600 nm for Calcein and 600 – 700 nm for Propidium Iodide. The 3D image was reconstructed (ZEN, Confocal microscope software) from slices acquired in a Z-stack mode with a plane interval of 1,5 μm.

### Immunostaining, labelling and Confocal Microscopy (Confocal Imaging Labelling)

Cells in the microfluidic chip were washed with 1x PBS (Gibco) and fixed with 4wt% solution of paraformaldehyde (PFA, Sigma Aldrich) in 1x PBS. After 10 minutes the fixative was washed away with 1x PBS, cells were permeabilized during 10 minutes with 0.1% solution of Triton X-100 (Sigma Aldrich) in 1x PBS and exposed for 1h to a 3% Bovine Serum Albumin (BSA, Sigma Aldrich) blocking solution in 1x PBS.

Next, the HUVECs’ tight junctions were stained using 5 μg/mL ZO-1 (Zonula Occludens-1) Monoclonal Antibody conjugated with Alexa Fluor 488 (Thermo Fisher Scientific) solution in previously prepared 3% BSA during O/N incubation at 4°C. In the next step the cells were washed with 3% BSA solution and incubated with 1x Phalloidin-iFluor594 (Abcam, stock 1000x) solution (in 1% BSA) for 30 min. at RT to stain actin filaments. The cell nuclei were stained after washing the cells with 1x PBS, using Hoechst 33258 stain at concentration 5 μg/mL. After 10 min. of incubation at RT the cells were washed with 1x PBS and imaged at RT using Zeiss LSM 800 Confocal microscope. Nuclei, tight junctions and actin were excited using 405 nm, 488 nm and 561 nm laser, respectively. The 3D images were acquired scanning the sample in a Z-stack mode, with an acquisition plane each 1 to 10 μm and later reconstructed into 3D image using the ZEN (Confocal microscope) software.

## Results and Discussion

### The system: amphiphilic PEG-dendron micelles and microfluidic cancer-on-a-chip

Three amphiphilic PEG-dendron hybrids, differing by their lipophilic end-groups in the hydrophobic block, were synthesized following a previously reported methodology,[34], [35] structures are shown in Figure 1a. The amphiphiles were labeled with a responsive dye, 7-diethylamino-3-carboxy coumarin, which forms and excimer when the hybrids self-assemble into micelles. The excimer formation causes a red-shift of the emission spectra of the dye allowing us to discriminate the assembled state (micelle) from the disassembled state (monomers), as shown in Figure 1a. The responsive properties of these micelles in serum and cells, have been previously demonstrated.[35] Here we aim to exploit these spectrally active micelles as a model system to study stability in a complex cancer-on-a-chip platform.

**Figure 1.**
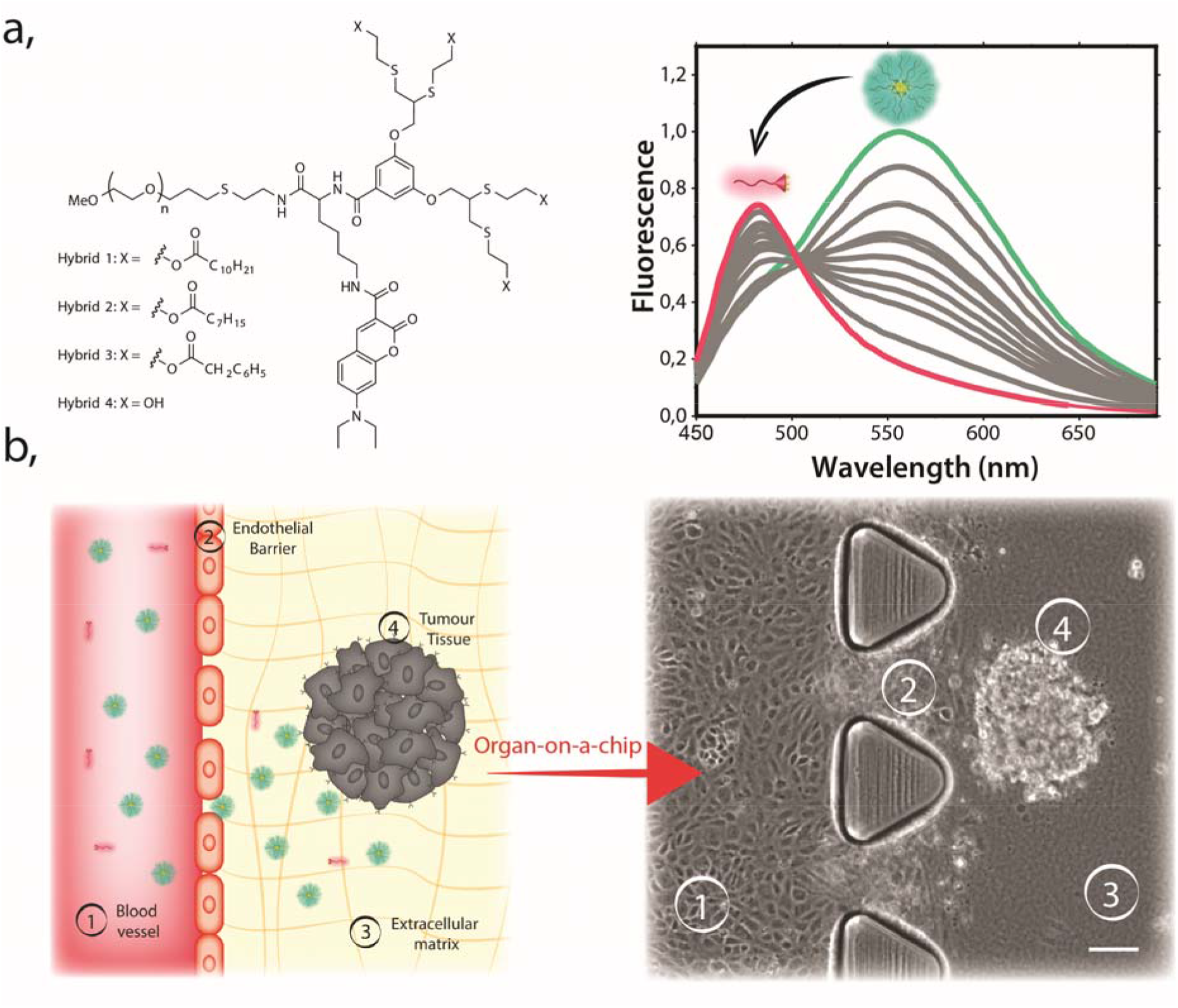
Schematic representation of our model. **a,** Molecular structure of the amphiphilic PEG–dendron hybrid polymers (left). Fluorescence emission graph for micelle-monomer equilibrium (right). In magenta monomer (fully disassembled structure) and in green fully assembled micelle. **b**, Schematic representation of the Cancer blood vessel-on-a-chip system, with marked the reconstructed barriers. Scale 100 μm.

With the microfluidic cancer-on-a-chip we aimed to recapitulate the four important barriers that the micelles will have to overpass when injected into the body: 1) the blood vessel circulation; 2) the endothelial barrier; 3) the ECM, and 4) the tumor spheroid (Figure 1b). To establish our platform and recreate these environments we used 3D cell culture chip[36], consisting of 3 microfluidic channels: the central channel with 1.3 x 0.25 mm (w x h) and the two media channels with 0.5 x 0.25 mm (w x h) dimensions (see Supporting Figure S1). The middle channel is separated from the two lateral channels by rows of triangular posts distant by 100 μm from one another, as shown in Figure 1b right. In our model, one of the lateral channels represents the blood vessel and the interface between the pillars reconstitutes the endothelial barrier. Human Umbilical Vein Endothelial Cells (HUVECs) were seeded on the upper and lower plane of the channel which induced the formation of a lumen-like geometry. The created EB separates the inner lumen of the blood vessel from the central microchannel, where we introduced preformed HeLa spheroids embedded into the Type 1 Collagen Gel. Importantly, coculture of endothelial and cancer cells in the same systems adds complexity to the model, therefore, growth kinetics of HeLa cells and HUVEC cells were previously evaluated to determine the optimal medium for the healthy growth of both cell lines (see Supporting Figure S2). Overall, we recreated a cancer-on-a-chip microenvironment where micelles stability can be evaluated during perfusion through the blood vessel channel. Moreover, the geometry of our model, where one channel is parallel to the other, enables continuous imaging of the interactions of nanosystems with all barriers. This real time imaging cannot be done in other channels conformations proposed previously in which only end time points studies could be performed.[33]

### Reconstructing the barriers in microfluidic cancer-on-a-chip

The evaluation of supramolecular stability using organ-on-a-chip requires a thorough design and characterization of the 3D model prior to further studies. Therefore, we performed a series of experiments aiming to validate the integrity and functionality of our chip. Figures 2a-c show transmission images of our model focusing on the two different channels with Figure 2a zooming into the vessel channel and Figure 2c into the gel embedded cancer spheroid, while Figure 2b shows a zoom-out image of the two channels.

**Figure 2.**
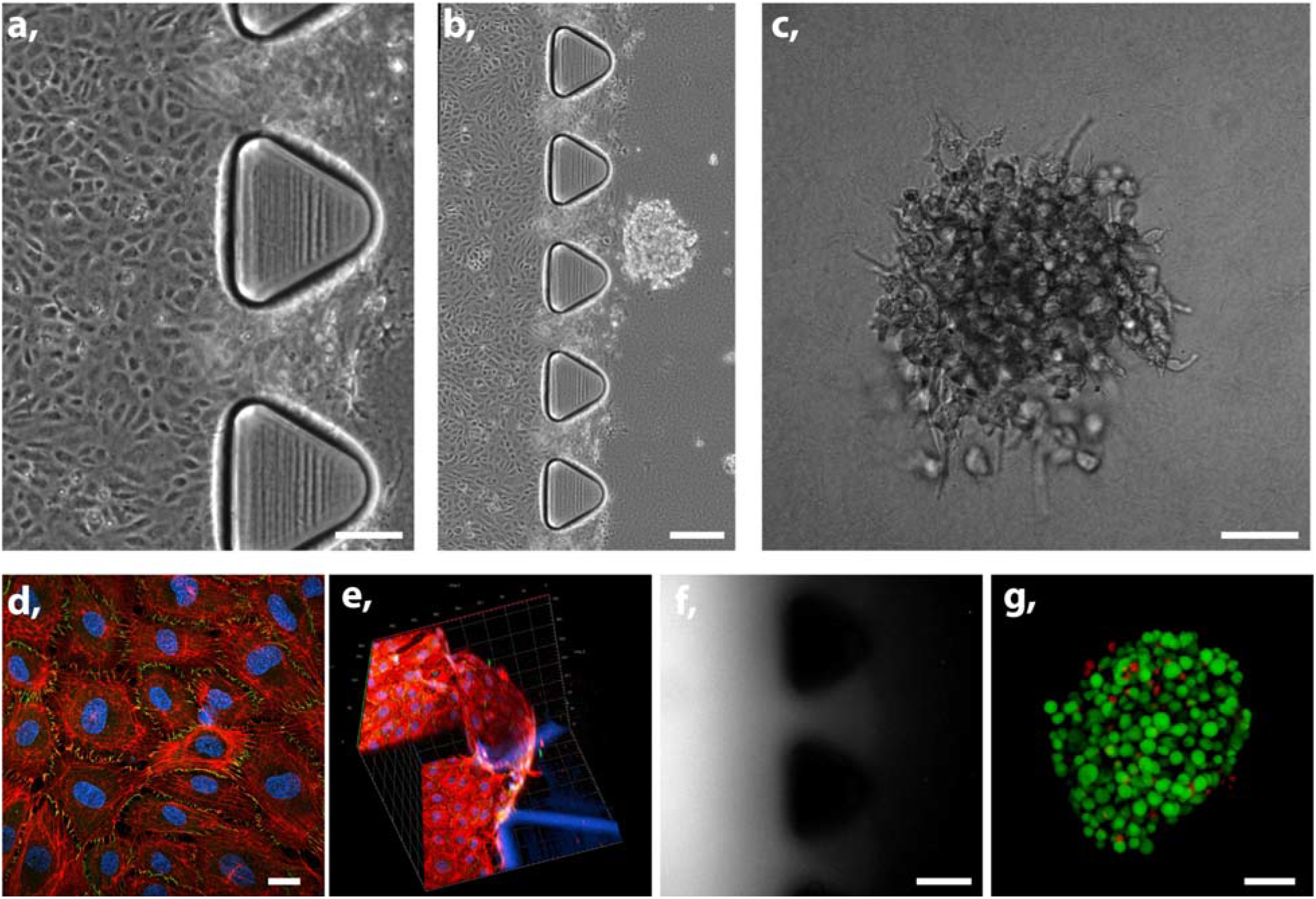
Tumor vessel-on-a-chip model. a) Transmission image of lateral channel zoomed in showing HUVEC monolayer. Scale bar 100 μm, b) Transmission image showing central and lateral channel of the chip with HUVEC in lateral and HeLa in central channel. Scale bar 200 μm, c) Transmission image zoomed in HeLa spheroid in the collagen gel. Scale bar 50 μm, d) Confocal image of HUVEC confluent monolayer (red: Phalloidin, blue: Hoechst, green: ZO-1). Scale bar 20 μm, e) 3D confocal image of HUVECs/gel interface (red: Phalloidin, blue: Hoechst, green: ZO-1), scale: axis ticks separation 40 μm, f) Epi fluorescence image of Dextran perfused during 10 minutes into the lateral channel lined with HUVEC monolayer, retaining the 10 kDa Dextran. In this experiment there were no HeLa cells in the collagen gel. Scale bar 150 μm, g) Confocal image of stained HeLa spheroid live – green (calcein), dead – red (Propidium iodide). Scale bar 50 μm

To validate our model, first we aimed to characterize the formation of an endothelial barrier separating both channels. The magnified transmission image of the vessel channel (Figure 2a) demonstrates the arrangement of a HUVEC monolayer after 3 days of unidirectional medium perfusion, which provided nutrients in a dynamic manner, mimicking physiological conditions, in both the lower and the upper plane of the channel (Supporting Figure S3 and S4). The formation of the confluent endothelial monolayer was further validated by fixating and staining the cells as shown in confocal image in Figure 2d, where actin, nucleus and formation of tight junctions can be observed. The endothelial cell to cell contact results in the expression of tight junction zonula occludens-1 (ZO-1, green) protein, which is essential in forming these junctions and corroborate the creation of a healthy endothelial barrier. Moreover, 3D confocal imaging allowed us to demonstrate the appearance of the EB on the interface of the blood vessel and ECM channels (Figure 2e). This vertical endothelial monolayer physically separates the lumen of the vessel channel from the collagen gel, mimicking the *in vivo* barrier. Interestingly, HUVECs in our chip aligned parallel to the flow direction due to the shear stress mimicking physiological conditions[37], [38](Supporting Figure S5). In contrast, cells cultured in static conditions show random arrangement following similar observations reported previously.[39] Subsequently, we tested the structural integrity of the HUVECs barrier by measuring the retention of fluorescently labeled 10 kDa Dextran perfused through the “blood vessel”. The fluorescence signal was detected in the lumen of the lateral channel but not in the collagen gel (Figure 2f) indicating proper functionality of the endothelial barrier. Altogether, these measurements indicate the formation of a functional EB with good structural integrity.

Finally, the transmission image of gel-embedded HeLa cells allowed us to observe their 3D spheroid conformation (Figure 2c). This arrangement implies less available surface area per cell, than in 2D cell culture models[40] and it recapitulates important aspects of geometry present in physiological conditions that cannot be studied in 2D cultures. The spheroids were introduced into the chip with sizes of 50-100 μm, and after 3 days of culturing they grew up 2-3 times of their original size, as shown in supporting Figure S3. A live/dead staining assay revealed that the cells are viable within the spheroid after 3 days of culture (Figure 2g), indicating that the nutrients of the culture medium and oxygen diffuse through the spheroids and are able to sustain the cell growth. Overall, we recreated a functional, microfluidic platform of a 3D tumor microenvironment, with perfusable blood vessel and cancer cells conformed into spheroid, improving the resemblance for the physiological conditions relevant for drug delivery compared to existing models.[29], [30], [32], [33]

### Increased extravasation of micelles is induced in tumor blood vessel chip

Having established our cancer vessel-on-a-chip microfluidic model, we aimed to investigate the ability of the micelles to permeate the blood vessel into the ECM channel. Previous studies using microfluidic models reported enhanced permeability of endothelial cells when exposed to specific molecules such as TNF-α[33] or when co-cultured with cancer cells,[30] leading to the formation of “leaky vessels” which recapitulate the EPR effect. However, the majority of these microfluidics models were used for studying only the end time point of nanoparticles permeation, lacking the intermediate progress of their extravasation. Herein, taking advantage of the design of our chip, we could continuously monitor the perfusion of our micelle system under three different experimental conditions: i) no HUVECs as negative control; ii) HUVECs barrier, and iii) HUVECs barrier with proximate HeLa spheroids mimicking a tumor blood vessel.

First, we aimed to test the ability of the micelle to cross the endothelial barrier, therefore, we perfused hybrid **1** into the three different models. In Figure 3a it can be observed that the hybrid immediately penetrates the collagen gel in the negative control (without HUVEC). This experiment proves that the properties of our NPs (*e.g.* small size of the micelles of about 20 nm[35]) allow their free diffusion through the ECM[5]. In contrast, hybrid **1** was successfully retained in the healthy blood vessel channel where HUVEC barrier blocks the permeation. Finally, if cancer cells were cultured in the collagen gel (mimicking the tumor microenvironment (TME)), we observed diffusion of hybrid **1** through the EB into the ECM channel after few minutes of infusion. These observations resemble experiments performed by Tang and coworkers where co-culture of cancer endothelial cells and breast cancer cells increased the permeation of nanoparticles through the EB.[30] Using quantitative analysis, we could define the percentage of polymer able to cross the barrier in each model as a function of the detected fluorescence intensity in the perfused channel as presented in the Figure 3c. We observed that close to 100% of the labeled hybrid fluorescence was detected in the collagen gel after less than a minute when there was no endothelial barrier. However, within the first 30 minutes of hybrid perfusion only an equivalent of 10% of the perfused polymer intensity was detected across the endothelial wall in the case of the healthy blood vessel, while at the same time about 80% of the signal of the hybrid in the vessel was detected in the ECM region in the tumor blood vessel model. These results indicate that the HUVEC monolayer in the healthy model significantly restricts the hybrid from crossing the barrier and penetrating the depth of ECM, however the barrier function was affected when HeLa spheroids are present in the gel (tumor model), where only a fraction of the perfused hybrid was retained by the endothelial wall. Interestingly, we hypothesized that the 10% of hybrid detected in collagen gel of the healthy model could origin from the permeation of the smaller monomer form (~3-5 nm) and not the assembled micelle (~20 nm), considering the healthy EB permeability.[41]

**Figure 3.**
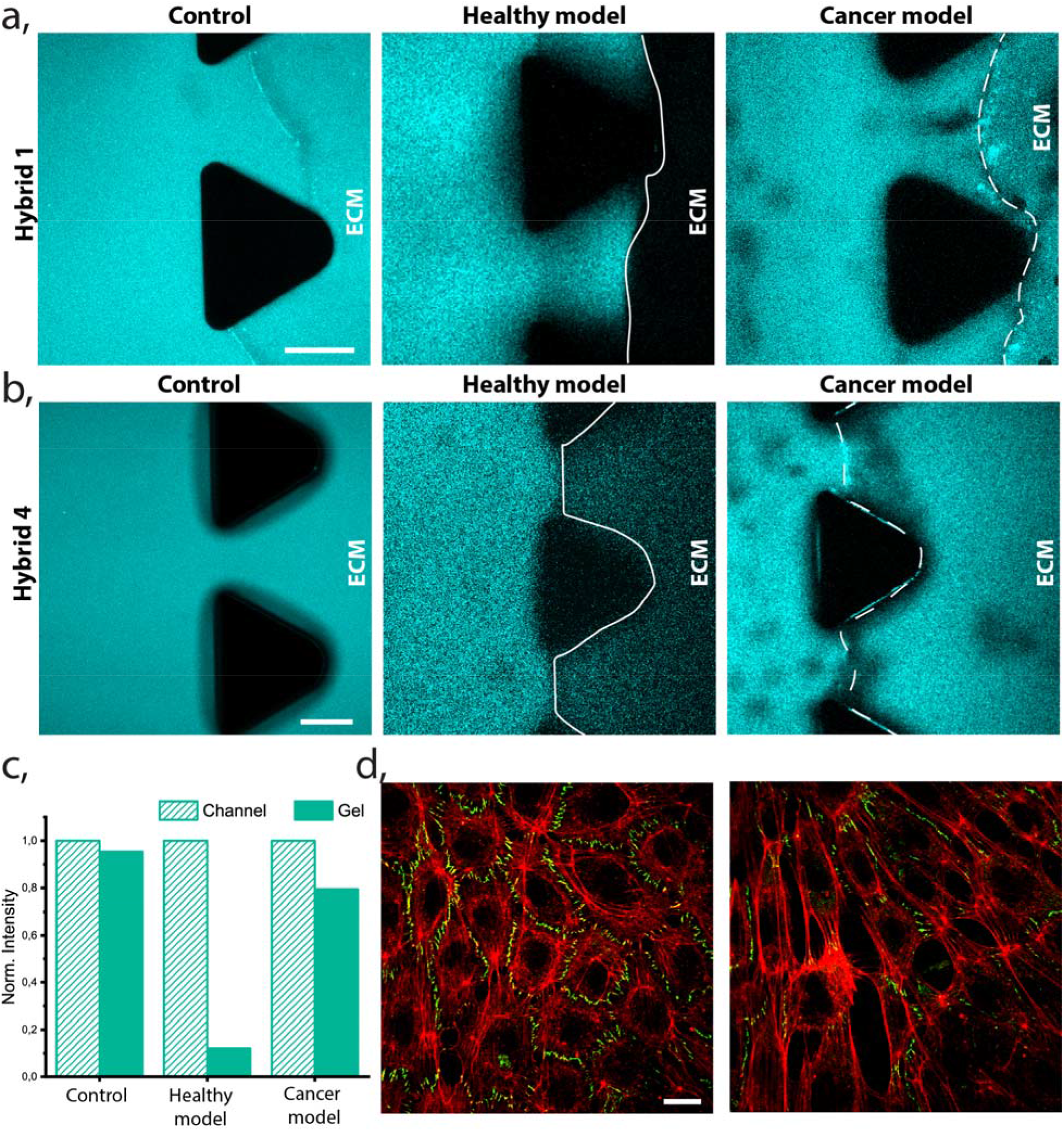
Extravasation of PEG-dendron hybrids in healthy and cancer models. **a.** Confocal image of hybrid 1 extravasation in a control chip (no HUVEC barrier), a healthy model (HUVEC barrier formed in the interface of the collagen gel) and cancer model (HUVEC barrier formed with presence of HeLa spheroids in the collagen gel). The images show emitted intensity between 446-700 nm, which include both monomer and micelle signal. Scale bar 100 μm **b.** Confocal image of hybrid 4 extravasation in the 3 corresponding models. Scale bar 100 μm **c.** Quantification of the average signal intensity of the hybrid 1 (Figure 3a) in the channel region compared to the gel region. **d.** Monolayer of HUVECs formed lining the healthy (left) and cancer (right) blood vessel channel. Actin (red) and ZO-1 (green). Scale bar 20 μm.

To investigate this proposed possibility of the monomer form to extravasate in the healthy model we perfused hydrophilic hybrid **4**, which has four hydroxyl end-groups and hence doesn’t self-assemble into micelles, as reported previously.[35] This hybrid was used as a control to study the behavior of the labeled monomeric form. The perfusion of hybrid **4** was followed in time in the three different models and the experiment was ended when the hybrid was observed to infiltrate into the collagen gel (like in Figure 3b). As expected, the monomer penetrates into the ECM in the control and the cancer models, but also, in the healthy one. HUVEC barrier partially retained the monomer in healthy blood vessel model for 25 minutes of continuous perfusion, limiting its concentration in the gel to ~40% of its concentration in the blood vessel channel. It is worth noting that after 5 minutes some hybrid could already be detected in the gel of healthy blood vessel (Supporting Figure S6). However, the time required was significantly shorter when HeLa cells were co-cultured in the gel. In case of tumor blood vessel, the penetration of hybrid **4** was almost immediate after inflow started, we measured similar concentrations of the hybrid in all regions after 5 minutes. Therefore, we hypothesize that the monomer form can cross into the ECM even in the healthy blood vessel model due to its small size, which may allow it to penetrate the ECM region through paracellular transport.

Next, we aimed to visually determine the morphological effect of co-culturing cancer cells on the structural integrity of the endothelial monolayer, to understand if a loss of integrity was the reason for the increased micelles permeation in the tumor blood vessel model. Tight junctions and adherent junctions are crucial structural elements formed between endothelial cells and an essential part of the endothelial barrier, regulating paracellular diffusion. These junctions’ main function is to restrict the permeation of molecules bigger than ~2 nm[42] from extravasation outside blood vessels. To further confirm that the enhanced permeability of the barrier in our microfluidic model was induced by the presence of cancer cells, we studied the expression of ZO-1, an essential protein forming tight junctions. Both, healthy and cancer blood vessel models were prepared as mentioned previously. After 3 days of continuous cell culture medium flow the cells were fixed in the chip and ZO-1 protein stained as shown in Figure 3d and supporting Figure S7. ZO-1 was clearly and uniformly expressed in the interface between HUVECs of our healthy model. However, the expression of ZO-1 was reduced when HeLa spheroids were cocultured with endothelial cells (cancer blood vessel-on-a-chip model). These observations indicated that cancer cells impact the HUVEC cell-cell interaction, reducing the tight junction formation, therefore, making the blood vessels leakier. All these observations further confirm the existence of an enhanced permeability in our cancer models. Similarly to previous studies by Kaji *et al*.[43] reporting that HUVEC and HeLa coculture affects the endothelial cells growth through either direct cell-cell contact as well as transmission of information via culture medium (paracrine communication). The cytokines excreted by HeLa can repulse HUVECs from their proximity and released reactive oxygen species lead to malfunction and even death of HUVECs, thus inducing leakiness of the endothelial barrier, permitting transportation of normally retained particles. However, it is worth noting that the permeation of micelles in the tumor blood vessel model was heterogenous and not always occurred at the same time point. We observed that small differences in number of spheroids or the proximity between spheroid and HUVECs monolayer impairs their retention capacity, which could be due to a variable concentration of signaling molecules (see Supporting Discussion). These observations reflect the heterogeneity of the EPR effect, which has been extensively discussed recently and is related not only to the type of cancer but also to the stage of the diseases.[44]

### Time- and space-resolved micelle stability revealed in 3D tumor microenvironment model

The major aim of the current work was to study the stability of our micellar systems when introduced into our microfluidic based 3D TME model. In our previous work, it was possible to detect the micelles and monomer inside cells due to the ability of the labeling coumarin dyes to change their emission wavelength upon disassembly. This self-reporting capabilities allow us to use confocal fluorescence microscopy to study the internalization pathway of different hybrids and correlate to them to their assembled state.[35] Herein, we hypothesized that the added complexity and dynamicity of the vessel-on-a-chip model may induce premature disassembly due to multiple possible interactions. To evaluate these critical interactions, we perfused micelles of hybrid 1, which was previously reported as the most stable system, in culture medium into the blood vessel channel. Then, we monitored in real time the micelles’ stability in key regions: the “blood vessel”, the endothelial barrier, the ECM and the HeLa spheroids as shown in Figure 4. Notably, micelles are represented in cyan and monomers (disassembled form) in magenta.

**Figure 4.**
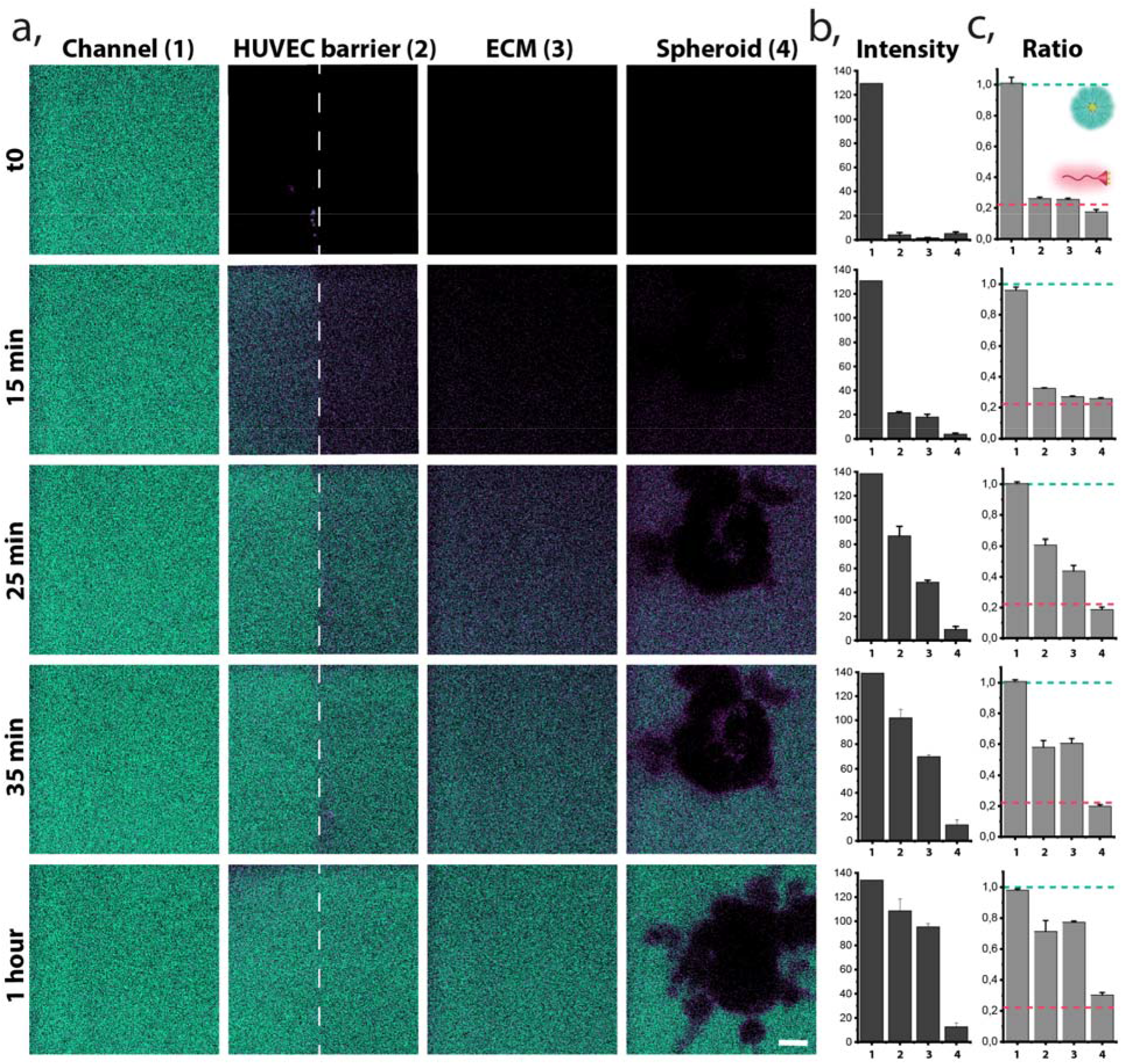
Space and time-resolved stability of hybrid 1. **a.** Ratiometric confocal images of real-time monitored presence of micelle (green) and monomer (magenta) at reconstructed barriers (1 - blood vessel, 2 - HUVECs barrier, 3 - ECM and 4 - cancer Spheroid) during constant perfusion of the hybrid 1, scale bar 15 μm. **b.** Time-resolved intensity of fluorescence signal (A.U) originating from the sum up of both micelle and monomer channels at each barrier. **c.** Normalized ratio of fluorescence signal between micellar and monomer form monitored in time at different barriers. Green dashed line indicates the ratio of fully formed micelles in equilibrium and the magenta dashed line indicates the ratio of fully disassembled (monomer) form.

In the first minutes of perfusion fluorescence was detectable only in the lateral perfused channel, coated with the monolayer of HUVECs (Figures 4a and b), and indicated assembled micelles (Figure 4c). Progression of hybrid penetration into the depth of the ECM was observed over time. After 15 minutes of continuous flow the hybrid started to reach the endothelial barrier, but based on the fluorescent intensity at the wave length corresponding to the monomeric coumarin, we could observe that mostly the monomer prevailed in passing through the ECs wall and entering the ECM as shown by the prevalence of the magenta signal in the central compartment. This is likely related to the smaller size of the monomer, allowing much better passage through the EB. After 25 minutes we observed the assembled micelles starting to traverse the EB, while the deep penetration into ECM (towards the tumor area) was still achieved mostly by the disassembled polymers. This observation could be attributed to two factors: i) the micelles progressively overcame the endothelial barrier or ii) the monomer form, which entered the ECM through the EB previously, accumulated and reached the critical micelle concentration (CMC), thus re-assembling into micelles. Figure 4b demonstrates that both, micelle and monomer forms coexist in the endothelial barrier, and in the ECM as the mean ratio is 0.6 and 0.4 respectively, not corresponding to the completely assembled neither the completely disassembled form. The assembled structures were not detected in the surroundings of the spheroids until more than half an hour of continuous perfusion. In addition, we observed only a weak penetration of the hybrid **1** into the depth of HeLa cells spheroids after 1 hour, but also after 2 hours of constant perfusion of micelles through the cancer blood vessel, as presented in Supplementary Figures S8 and S9. We observed the stabilization monomer/micelle equilibrium in all monitored regions after 1h from the beginning of the perfusion, excluding the spheroid. Interestingly, hybrid **1** in contact or in close proximity to HeLa cells was always detected as monomer. This observation contrasts to previously reported internalization behavior in 2D cell cultures where it was demonstrated that assembled hybrid **1** was taken up by HeLa cell via endocytosis, and its disassembly progressed in time.[35] This discrepancy can be attributed to the 3D conformation of the cells, as spheroids, which could promote different endocytosis processes related to the new 3D cells confluency and conformation, highlighting the importance of going beyond 2D cell culture models. Other preceding works investigated the penetration of nanosystems into tumor spheroids as a function of nanocarrier size, shape, charge and functionalization.[31], [45], [46] Likewise, the penetration of cross-linked and non-cross-linked micelles into spheroids has been compared, showing an improved tumor penetration for the cross-linked ones, which was linked with their higher stability.^48–50^ Therefore, we hypothesized that the lower penetration into our model can be caused by a premature disassembly in the periphery of the spheroid.

Here, for the first time, we can demonstrate the real-time interactions of dynamic supramolecular systems with cells in a dynamic 3D environment, showing significant differences than in a static 2D cell cultures. Therefore, the system stability at different physiological barriers can be evaluated and correlated to its final penetration into the cancer cells. These results highlight the importance of selecting adequate screening models, which resemble to much greater extent the *in vivo* cell arrangement and features, to rationally evaluate and optimize supramolecular nanocarriers design. This approach sheds light on how nanocarriers stability affects their accumulation in solid tumors *in vivo*. Overall, by combining confocal imaging and a microfluidic 3D cancer-on-a-chip model we could demonstrate the high stability of hybrid **1** against induced flow shear stress and interactions with the ECM, while we prove a low stability/penetration in contact with 3D arranged cancer cells. This impedes the assembled state to penetrate the solid tumor in the current more realistic model along with general low penetration of the perfused structures into the HeLa spheroids.

### Stability of hybrids dictates their infiltration/extravasation

Finally, we aimed to investigate the interplay between molecular structure and micellar stability and the consequent ability to extravasate. Therefore, we compared the stability of three hybrids, with decreasing length of the hydrophobic end-groups starting from four undecanoate tails for hybrid **1** to four octanoate alkyl chains for hybrid **2** and four phenyl acetate for hybrid **3**, which have also eight carbons but differ by the presence of the more polar aromatic ring. Using the cancer vessel-on-a-chip model, we wanted to identify the factors affecting their performance in the new 3D system. In our previous studies,[35] we demonstrated that stabilities of hybrid **2** and **3** were similar when diluted with serum, however, the kinetics of their disassembly were significantly different. While hybrid **3** disassembles rapidly upon dilution, hybrid **2** needed hours to reach the equilibrium. In contrast, hybrid **1** was stable both in the presence of serum proteins and upon dilution. Here, we aimed to understand how these differences in thermodynamic and kinetic stability are reflected in a complex model, where the micelles face the reconstructed barriers of the tumor microenvironment. Experiments with each one of the three hybrid systems were performed by perfusing a solution of culture media containing the micelles through the blood vessel channel of the chip. In Figure 5 we show representative images of two different areas of the chip for each of the perfused micelles. First, we investigated the ability of each hybrid to cross the endothelial barrier. We observed extravasation of all hybrids **1-3** when HeLa spheroids were located close to HUVECs (Figure 5b). In contrast, in the regions where spheroids were further away (hundreds of microns), only hybrid **3**, which has the least hydrophobic dendron in comparison with hybrids **2** and **1**, was able to significantly extravasate to the ECM. Thus, we conclude the endothelial barrier “leakiness” can be heterogenous, depending on the amount, distance and distribution of tumor spheroids in the ECM.

**Figure 5.**
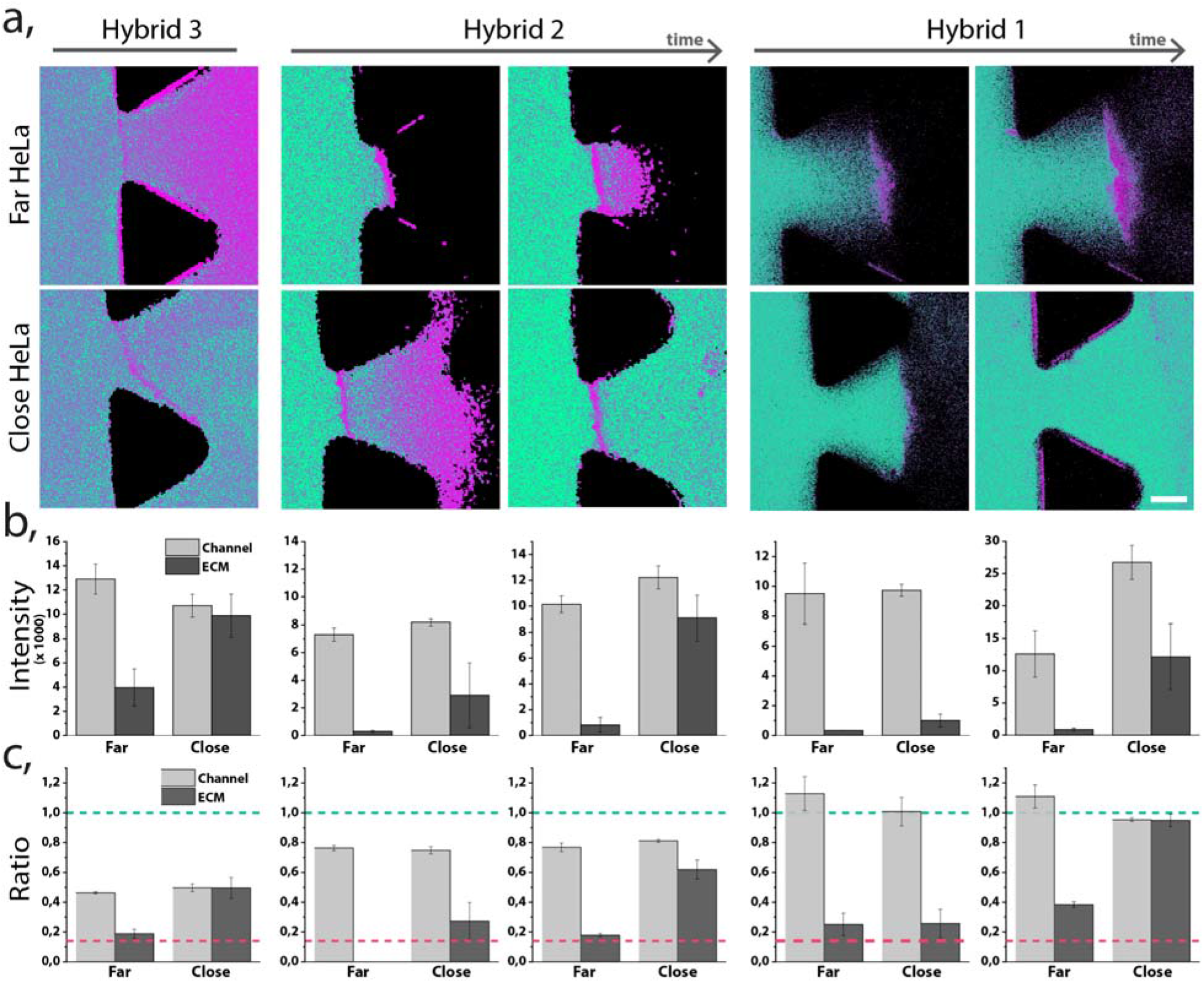
Stability of different hybrids in blood vessel-on-a-chip. **a.** Ratiometric confocal images of the different hybrids inside the chip in two different regions. Hybrid 2 and 3 are shown at 2 different time points: less than 15 minutes and after more than 30 minutes of continuous perfusion. Scale bar 75 μm. **b.** Intensity of fluorescence signal originating from the sum up of the intensity of both the monomer and the micelle channels. Intensity was measure in the vessel channel and in the ECM region **c.** Normalized ratio of fluorescence signal between micellar and monomer form for each hybrid and in each region. Green dashed line indicates the ratio of fully formed micelles in equilibrium and the magenta dashed line indicates the ratio of fully disassembled (monomer) form.

We next tested the stability of the three hybrids, under flow conditions when perfused through the vessel-like channel (Figure 5a and c). Based on the ratios between the emitted intensity of the coumarin dye at 480 nm and 550 nm, corresponding to the disassembled and assembled state of the labeled polymers, respectively, we could study the assembly states of the different hybrids. We did not observe disassembly of hybrid **1** in the vessel channel, with mean ratio of fluorescence signal between micellar and monomer form equal to 1, which is indicative of the presence of micelles, while we observed slight disassembly of hybrid **2** and significant disassembly of hybrid **3**, with mean ratios of 0.8 and 0.5 respectively. Previous investigations[35] showed hybrid **2** and **3** were slightly unstable in presence of serum proteins, but the degree of disassembly based on the fluorescent ratio was the same for both. Therefore, we hypothesized the enhanced disassembly rate of hybrid **3** is not only due to interactions with serum proteins but also affected by the flow. This result indicates that the flow-induced shear stress can drastically affect the stability of supramolecular nanocarriers.

Further, we monitored the stability of each hybrid at the previously defined barriers and observed significant differences among them. Interestingly, in regions far from spheroids only the monomeric form of hybrid **3** was able to efficiently extravasate. This phenomenon could occur due to the increased disassembly of the micelle in contact with the HUVEC barrier, which allowed the monomer i) to paracellularly extravasated due to its small size or ii) to transcellularly cross the EB. Noticeably, among all the tested hybrids only monomer of hybrid **3** extravasates efficiently, most likely due to the low stability of this hybrid in contact with cells. In contrast to that, in regions close to HeLa, hybrid **3** crossed the EB mostly in the semi-assembled state (as it appeared in the blood vessel channel), probably due to the increased wall leakiness. These results indicated that the disappearance of the tight junctions allowed the assembled micelles to cross the barrier. Differently, extravasation of micelles formed from hybrid **2** in regions close to HeLa, had a time dependent response; first only monomer crossed the EB whereas, assembled micelles were detected in the ECM only after more than 30 minutes. Finally, we observed that hybrid **1** behaves similarly to the hybrid **2**, where monomer molecules extravasated first, followed by the later penetration of the assembled micelles. Interestingly, while hybrid **2** and **3** accumulated in the endothelial barrier only as a monomer, hybrid **1** monomeric form accumulated only in areas far away from the HeLa cells. We hypothesize a relationship between the micelle stability and the different internalization behavior of hybrids, while **3** and **2** were observed to internalize as monomer in the previous work, **1** internalized as a micelle and disassembled over time[35]. These results corroborate our previous observations, where hybrid **1** is proven to be the most stable system, but also, demonstrate the high stability of hybrid **2** in complex systems. Overall, we could correlate the interplay between stability of the micelles and their performance in a 3D model, as well as their ability to extravasate to reach the cancer cell regions.

## Conclusions

In the present work, we introduced the combination of spectral confocal imaging and a microfluidic 3D cancer-on-a-chip model as a new approach to study the stability of supramolecular nanocarriers. The unique fluorescence properties of our micelles allowed us to track their assembly state across the changing conditions and correlate their stability with the ongoing biological interaction. The cancer-on-a-chip introduced here, enriches the study thanks to its similarities to physiological conditions, introduced especially via the 3D dimension in cellular distribution and through active flow in the blood vessel channel. This approach helps to recapitulate the barriers to be overcome (*e.g.* blood flow, endothelial wall, ECM and 3D cancer spheroids) that cannot be successfully reconstructed in a 2D cell culture. The results obtained show the formation of leaky vasculature in the presence of cancer cells, but also, a high heterogeneity among different chips or even across distant regions of the same chip, related to number of cancer cells and distance from the endothelial cells. Importantly, these features resemble to a great extent the *in vivo* pathologies of many tumors. Moreover, we obtained a precise and direct information about the performance and stability of the micelles in each of the barriers, thanks to the time and space-resolved imaging. We demonstrate the ability of hybrids **1** and **2** to extravasate from ‘blood vessel’ as assembled micelles, while the complex interactions of EB and the ECM induce the disassembly of micelles of hybrid **3**. Interestingly, we observed the loss of stability of hybrid **1** in close proximity to spheroids, as well as, a very low penetration into the tumor. These results demonstrated the importance of screening nanocarriers stability in more complex 3D based systems to accurately predict their potential to be translated into the clinic. Our approach of combining spectrally responsive supramolecular structures with a platform reconstructing cancer-on-a-chip has the capacity to provide new knowledge about nanoparticles performance, stability and accumulation in tumor, which will allow to bridge the gap between *in vitro* and *in vivo* testing of new drug delivery systems.

## ASSOCIATED CONTENT

### Supporting Information

The following files are available free of charge.

Supporting Figures and Discussion (PDF file)

## ACKNOWLEDGMENT

NFG and AGM thank to D. Izquierdo for the experimental support and scientific discussions. The authors thank the Biomaterials for Regenerative Therapies group in IBEC for sharing their equipment to execute the experiments and IBEC’s Microscopy Characterization Facilities equipped with Confocal Microscope. LA and SP thank the Spanish Ministry of Economy, Industry and Competitiveness, through the Project SAF2016-75241-R, by the Generalitat de Catalunya through the CERCA program. The authors also acknowledge the EuroNanoMed II platform through the project NANOVAX and the foundation Obra Social La Caixa and the European Research Council (ERC-StG-757397). JS and RRT have support from the CERCA Programme and by the Commission for Universities and Research of the Department of Innovation, Universities, and Enterprise of the Generalitat de Catalunya (2017 SGR 1079). This work was partially funded by the Spanish Ministry of Economy and Competitiveness (MINECO) through the projects MINDS (Proyectos I+D Excelencia + FEDER): TEC2015-70104-P, BIOBOT (Programa Explora Ciencia / Tecnología): TEC2015- 72718-EXP and EuUONANOMED II PCIN-2016-025. RJA thanks the ISRAEL SCIENCE FOUNDATION (grant No. 1553/18) for the support of this research. The project that gave rise to these results received the support of a fellowship from “la Caixa” Foundation (ID 1000010434) The fellowship code of AGM is LCF/BQ/DI17/11620054. This project has received funding from the European Union’s Horizon 2020 research and innovation programme under the Marie Skłodowska-Curie grant agreement No. 713673.

## ABBREVIATIONS

DDS: drug delivery systems
DMEM: Dulbecco’s Modified Eagle Medium
EB: endothelial barrier
ECs: Endothelial Cells
ECM: extracellular matrix
EPR: Endothelial Permeability and Retention effect
(FRET): Förster Resonance Energy Transfer
HUVEC: Human Umbilical Vein Endothelial Cells
NPs: nanoparticles
TME: Tumor Microenvironment

