## Supplementary Information for "Real-time Ratiometric Imaging of Micelles Assembly State in a Microfluidic Cancer-on-a-chip"

**Supporting Figures**

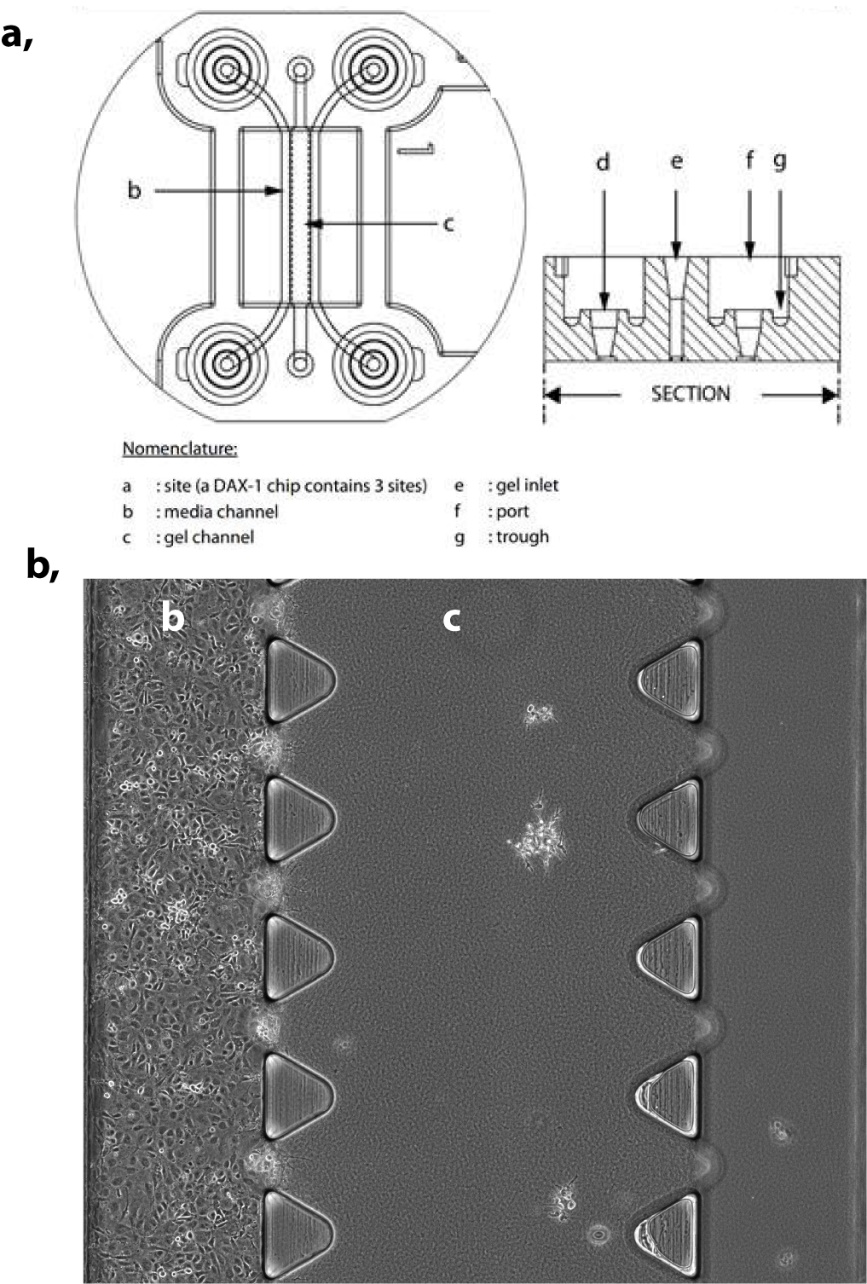

**Figure S1.** Microfluidic chip used to reproduce cancer-on-a-chip. It consists of three microfluidic channels the central channel with 1.3 x 0.25 mm (w x h) and the two media channels with 0.5 x 0.25 mm (w x h) dimensions. The middle channel is separated from the two lateral channels by rows of triangular posts distant by 100 µm from one another.

**Figure S2.** Growth kinetics of HUVECs and HeLa. **a.** HUVECs were seeded in a 96 well plate at a density of 2500 cells/well and incubated with three different medium types: DMEM medium supplemented with 10% FBS (which is used for HeLa cell monoculture) and Promocell or Millipore media optimized for HUVECs. PrestoBlue cell viability test was performed after 1, 3 and 4 days of incubation. The absorbance at 570 nm is plotted as a function of time. HUVECs did not grow in DMEM medium but they grew similarly using both HUVEC optimized media. **b.** HeLa cells were seeded at a density of 2500 cells/well in a 96 well plate and incubated with the two HUVEC media to decide which was the optimal for HeLa grow. PrestoBlue cells viability test was performed after 1, 3 and 4 days. The graph shows the absorbance intensity as a function of time. HeLa cells growth kinetic
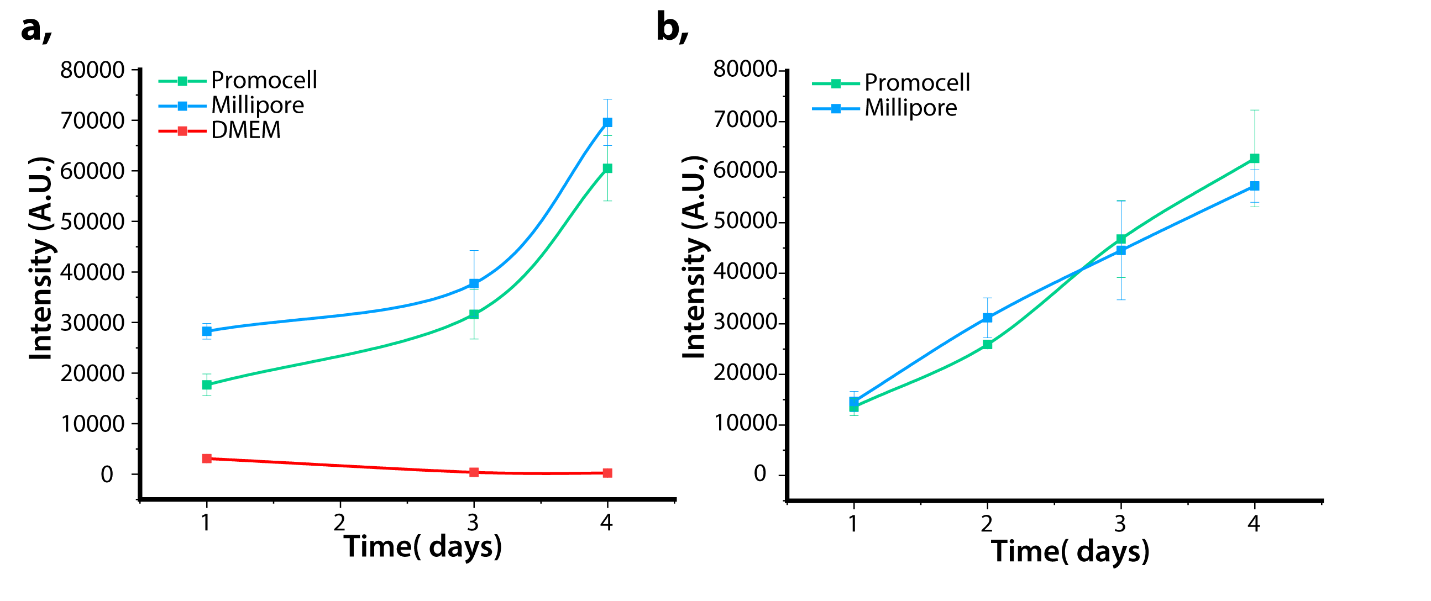
was similar using both media. Every condition was performed in sextuplicate, error bar represents S.D. between wells.

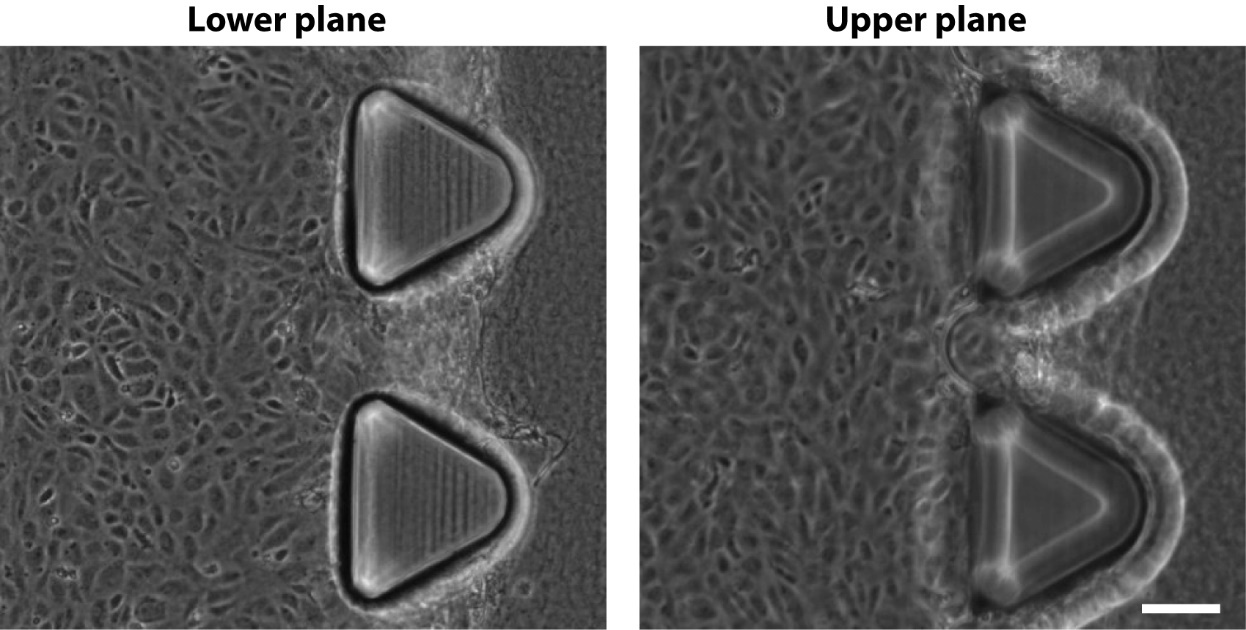

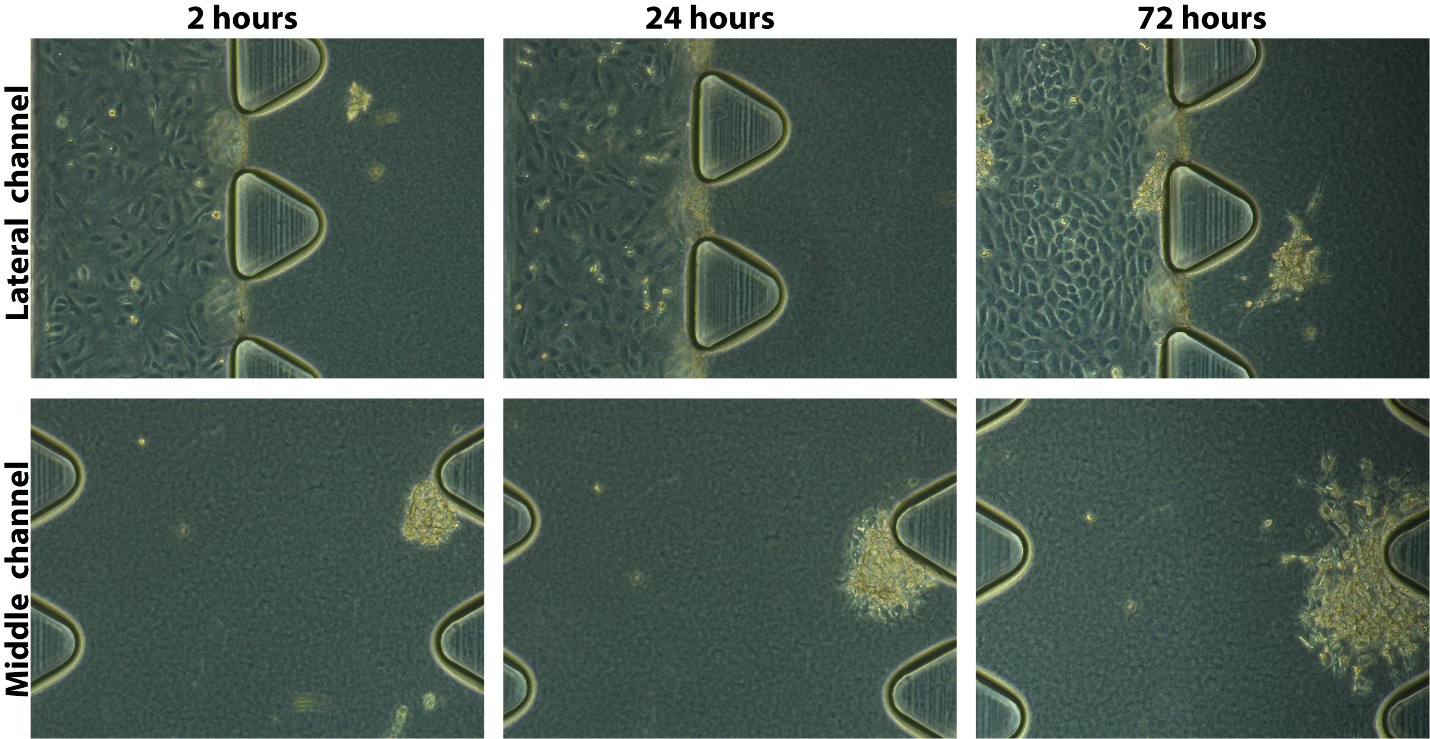
**Figure S3.** Cell growth in the microfluidic chip after 2 hours, 24 hours and 72 hours. Transmission images of the same chip are shown: upper row is a magnified image of the lateral channel were HUVECs were seeded to form the ‘blood vessel’ and part of the middle channel. Lower row shows magnified images of the middle channel consisting on the ECM and the HeLa spheroids. Distance between triangular posts is 100 µm.

**Figure S4.** Transmission image of the ‘blood vessel’ channel after 72 hours of continuous medium perfusion. Both the lower and the upper plane of the channel are shown to demonstrate the formation of a cell monolayer in both planes. Scale bar 100 µm.

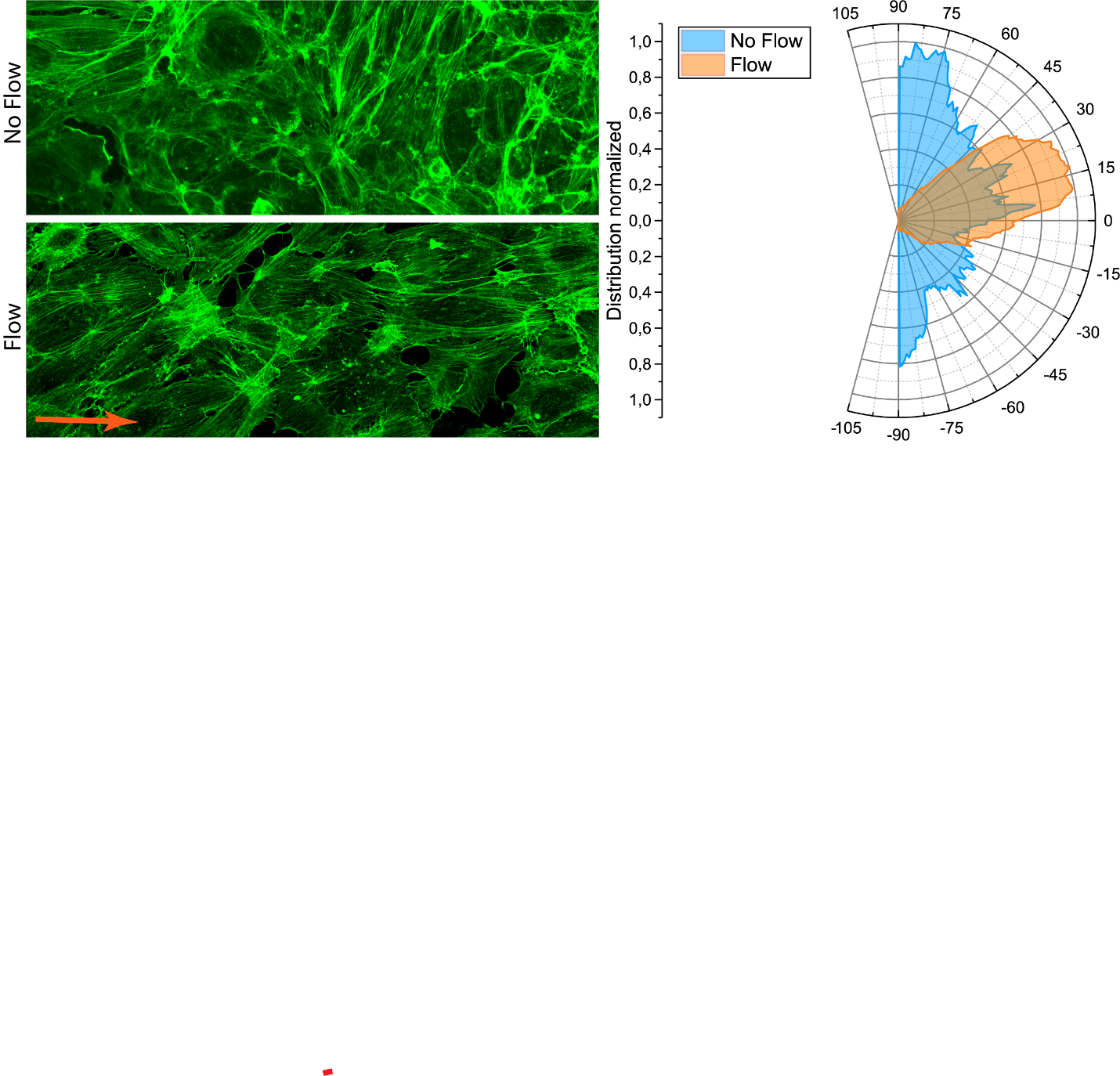

**Figure S5.** HUVECs alignment with the direction of the flow. Two independent chips were prepared as explained, however, one of the chips was incubated in static conditions, with medium change every 24h, meanwhile the other was continuously perfused with cell medium. After 72 hours the cells were fixed and actin stained, confocal images of actin were acquired using Zeiss LSM 800 microscope. The images were analyzed using the OrientationJ plugin of ImageJ to obtain the distribution of the orientation graphs. It can be observed that the cells without flow are oriented randomly having components in all directions, while the cells cultured under flow have a dominant direction coinciding with the medium flow.

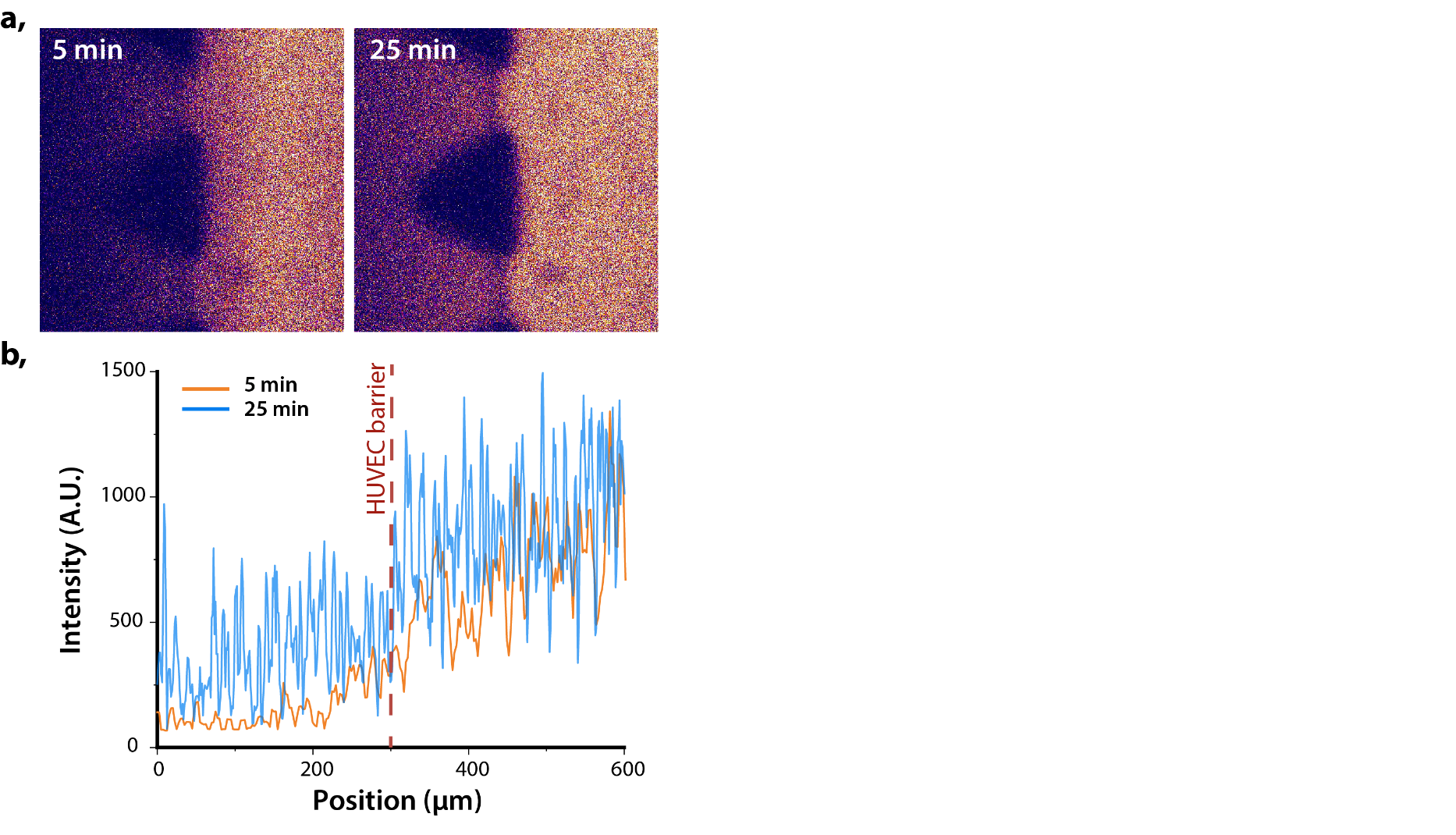

**Figure S6.** Extravasation of hybrid 4 in healthy model already occurs after 5 minutes of continuous perfusion. A healthy microfluidic model was prepared and hybrid 4 was flow for 25 minutes to study its extravasation **a.** Confocal images of the hybrid 4 being flow through the ‘blood vessel’ at 5 and 25 minutes. In the images the ‘blood vessel’ channel (right) and part of the ECM (left) can be observed. Look-up table “fire” is used for visualization purposes **b.** Plot profile of a horizontal line across each image showing the intensity detected in each position. After 5 minutes hybrid was detected in both: the gel and the lateral channel, however the intensity in the gel was only 15% comparing to the channel, while after 25 minutes it raised to 40%.

.
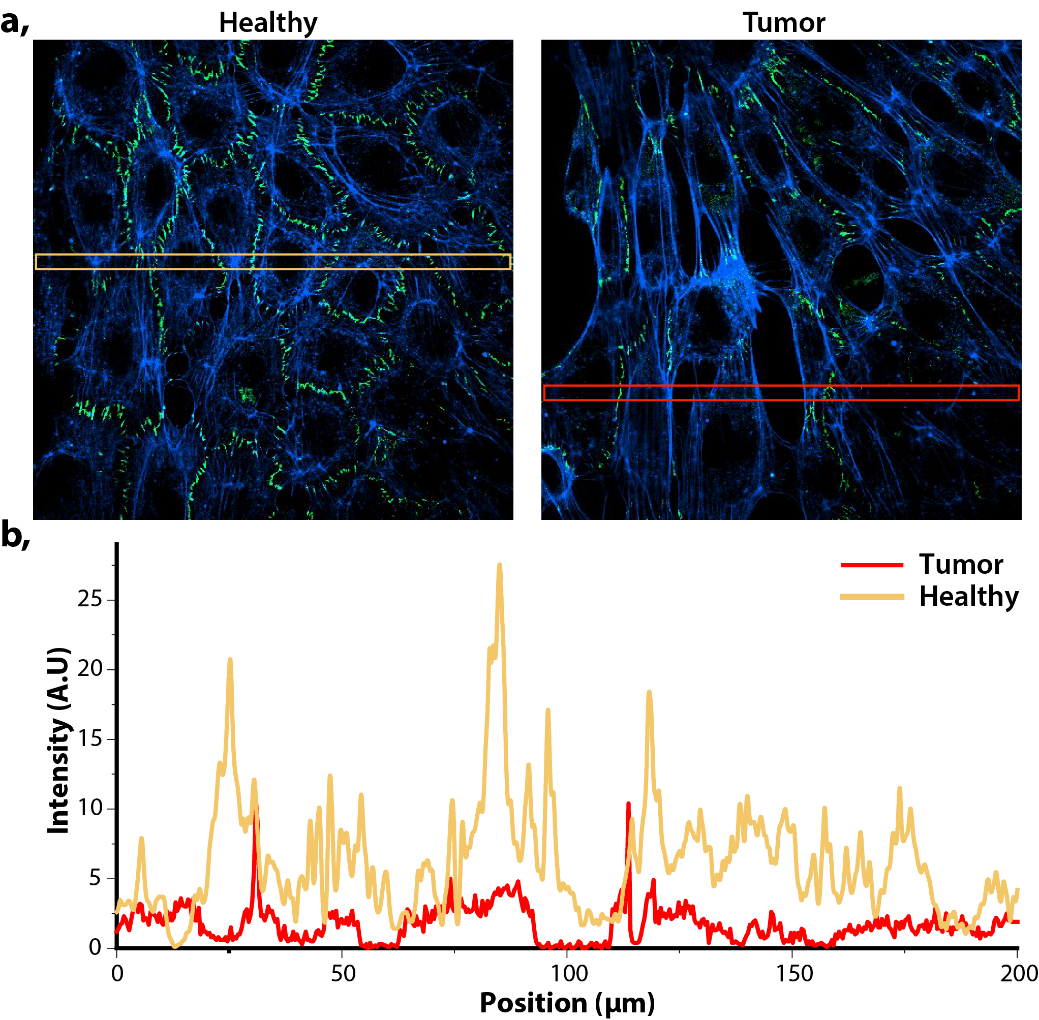
**Figure S7. a.** Confocal images of monolayer of HUVECs formed lining the healthy (left) and cancer (right) blood vessel channel. Actin (blue) and ZO-1 (green). **b.** Plot profile of ZO-1 expression of the section highlighted in (a) of cancer vs healthy model.

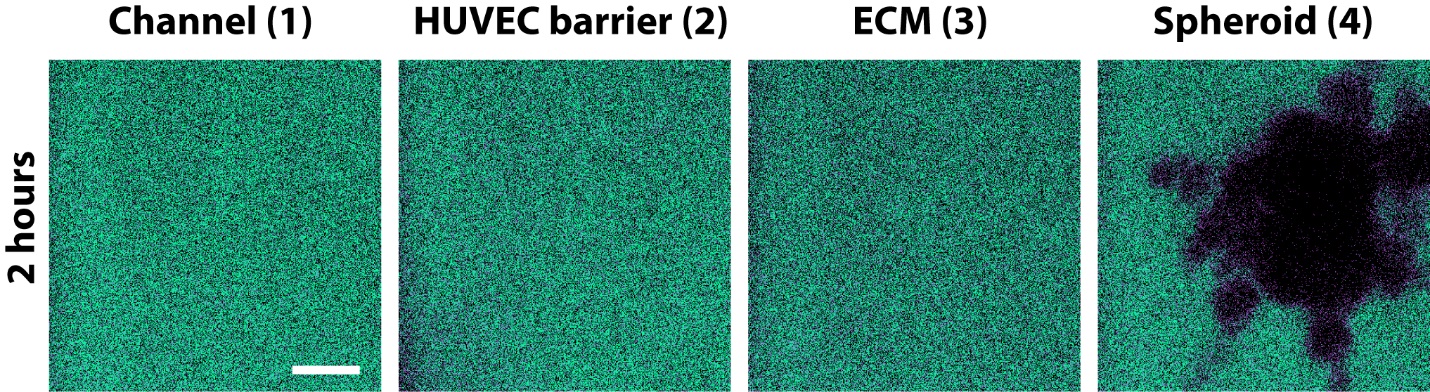

**Figure S8. S**pace resolved stability of hybrid **1** after 2 hours of continuous perfusion. Ratiometric confocal images of real-time monitored presence of micelle (green) and monomer (magenta) at reconstructed barriers (1 - blood vessel, 2 - HUVECs barrier, 3 - ECM and 4 - cancer Spheroid). Scale bar 20µm.

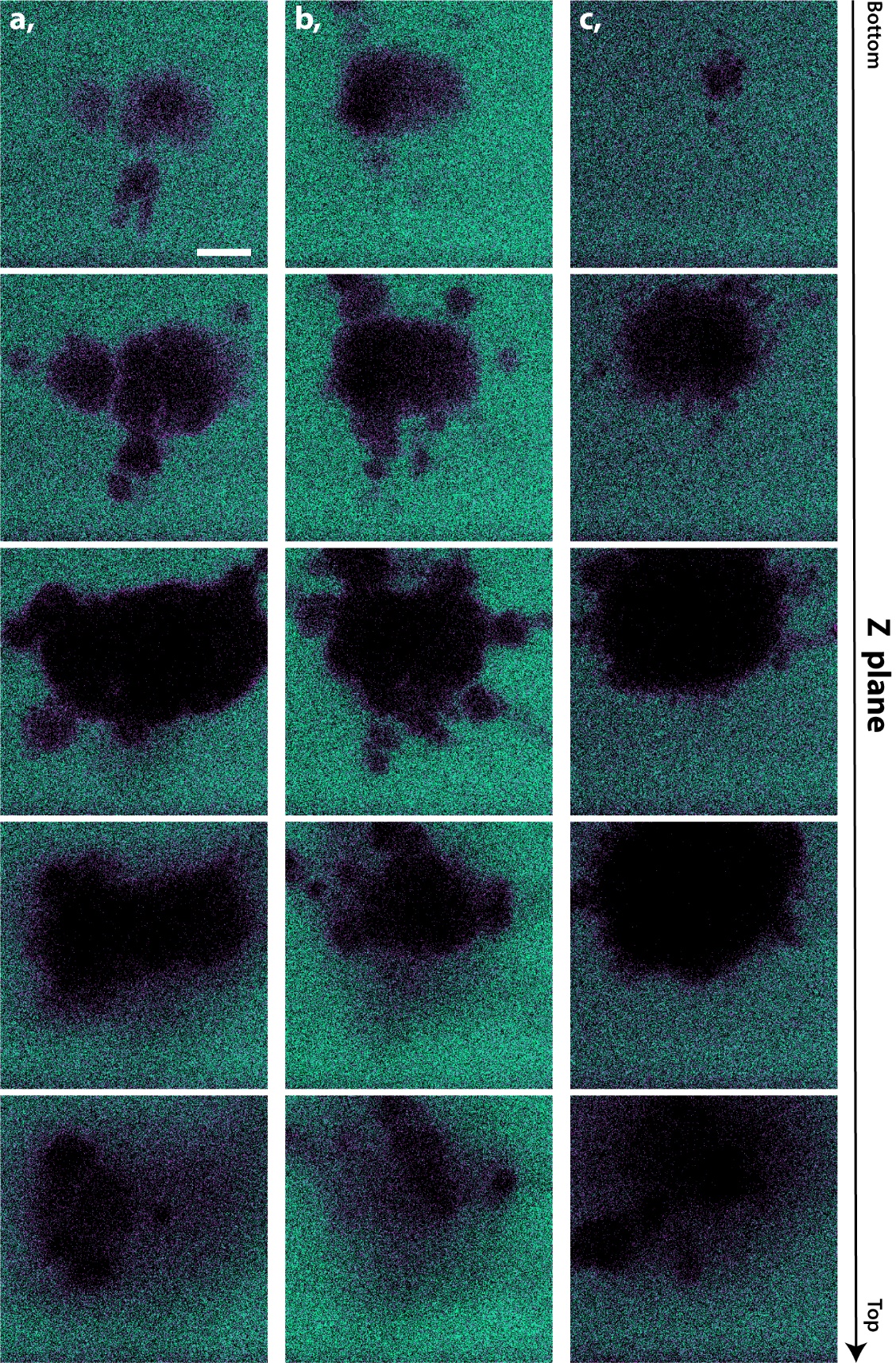
**Figure S9.** Penetration and stability of hybrid 1 in the HeLa spheroids after 2 hours of continoius perfusion. Ratiometric confocal images of real-time monitored presence of micelle (green) and monomer (magenta). **a.** Z-stack images of spheroid 1 **b.** Z-stack images of **s**pheroid 2 **c.** Z-stack images of **s**pheroid 3. Scale bar 20µm

**Supporting Discussion**

The preparation of tumor model chip consisted on a procedure described in the Materials and Methods section of the article. Despite the followed protocol was in all experiments the same, each chip preparation presented few differences, also reflected when infusing hybrids. Those differences relate mostly to the number, and distribution of spheroids within the extracellular matrix (collagen gel). Upon chip preparation we could adapt the number of spheroids from few to over a dozen and adjust the number accordingly through a visual inspection. However, the distribution of spheroids across the ECM can be hardly controlled, especially their distance from the endothelial barrier. Within the performed experiments (infusion of hybrids into the blood vessel lumen) we observed certain heterogeneity in the retention of perfused material in the HUVECs lined channel. We registered either immediate monomer/micelle crossing the EB or more gradual, depending on the regions of the microfluidic chip. Overall, the immediate passage of the perfused polymer was attributed to the HeLa spheroids present very closely to the EB (less than tens of micrometers), in such cases the hybrids entered the gel channel in less than a minute. On the other hand, when the spheroids were located further away (more than tens of micrometers) the HUVEC wall appeared leaky, but the rate of perfused structures entering the ECM was variable. Depending on the tested chip, the detected fluorescence intensity was similar on both sides of the HUVECs wall within few to tens of minutes and in some areas even after 30 minutes. As reported in literature, the coculture with HeLA affects the endothelial cells, not only through cell-cell contact (when we observed very rapid entering of the hybrids into the ECM), but also via paracrine communication. We hypothesize that this transmission of information via culture medium cannot be easily controlled in dynamic conditions and 3D spheroid random distribution, as it could be in a static setup, where the system is allowed to reach an equilibrium. As the HeLa-excreted molecules are harmful to HUVECs, their integrity becomes impaired and the effect likely depends on the exposure to these damaging species. Overall, this heterogenous leakiness of HUVEC barrier reflects the EPR heterogeneity observed *in vivo*, which is highly dependent on cancer type, stage and patient.
